# GRP78 and Integrins Play Different Roles in Host Cell Invasion During Mucormycosis

**DOI:** 10.1101/2020.04.29.069666

**Authors:** Abdullah Alqarihi, Teclegiorgis Gebremariam, Yiyou Gu, Marc Swidergall, Sondus Alkhazraji, Sameh S.M. Soliman, Vincent M. Bruno, John E. Edwards, Scott G. Filler, Priya Uppuluri, Ashraf S. Ibrahim

## Abstract

Mucormycosis, caused by *Rhizopus* species, is a life-threatening fungal infection that occurs in patients immunocompromised by diabetic ketoacidosis (DKA), cytotoxic chemotherapy, immunosuppressive therapy, hematologic malignancies or severe trauma. Inhaled *Rhizopus* spores cause pulmonary infections in patients with hematologic malignancies, while patients with DKA are much more prone to rhinoorbital/cerebral mucormycosis. Here we show that *R. delemar* interacts with glucose-regulated protein 78 (GRP78) on nasal epithelial cells via its spore coat protein CotH3 to invade and damage the nasal epithelial cell. Expression of the two proteins is significantly enhanced by high glucose, iron and ketone body levels (hallmark features of DKA), potentially leading to frequently lethal rhinoorbital/cerebral mucormycosis. In contrast, *R. delemar* CotH7 recognizes integrin β1 as a receptor on alveolar epithelial cells causing the activation of epidermal growth factor receptor (EGFR) leading to host cell invasion. Anti-integrin β1 antibodies inhibit *R. delemar* invasion of alveolar epithelial cells and protect mice from pulmonary mucormycosis. Our results show that *R. delemar* interacts with different mammalian receptors depending on the host cell type. Susceptibility of patients with DKA primarily to rhinoorbital/cerebral disease can be explained by host factors typically present in DKA and known to upregulate CotH3 and nasal GRP78 thereby trapping the fungal cells within the rhino-orbital milieu, leading to subsequent invasion and damage. Our studies highlight that mucormycosis pathogenesis can potentially be overcome by the development of novel customized therapies targeting niche-specific host receptors or their respective fungal ligands.

**Importance:** Mucormycosis caused by *Rhizopus* species is a fungal infection with often fatal prognosis. Inhalation of spores is the major route of entry, with nasal and alveolar epithelial cells among the first cells that encounter the fungi. In patients with hematologic malignancies or those undergoing cytotoxic chemotherapy, *Rhizopus* causes pulmonary infections. On the other hand, DKA patients predominantly suffer from rhinoorbital/cerebral mucormycosis. The reason for such disparity in disease types by the same fungus is not known. Here we show that, the unique susceptibility of DKA subjects to rhinoorbital/cerebral mucormycosis is likely due to specific interaction between nasal epithelial cell GRP78 and fungal CotH3, the expression of which increase in the presence of host factors present in DKA. In contrast, pulmonary mucormycosis is initiated via interaction of inhaled spores expressing CotH7 with integrin β1 receptor which activates EGFR to induce fungal invasion of host cells. These results introduce plausible explanation to disparate disease manifestations in DKA versus hematologic malignancy patients and provide a foundation for development of therapeutic interventions against these lethal forms of mucormycosis.

## Introduction

Mucormycosis is a lethal infection caused by mold belonging to the order Mucorales (1, 2). The infection is characterized by high degree of angioinvasion which results in substantial tissue necrosis, frequently mandating surgical debridement of infected tissues (3, 4). Despite aggressive treatment with surgical removal of infected foci and use of the limited options of antifungal agents, mucormycosis is associated with dismal mortality rates of 50-100% (5, 6). Also, surviving patients often require major reconstructive surgeries to manage the ensuing highly disfiguring defects (2, 7).

*Rhizopus* spp. are the most common etiologic agents of mucormycosis, responsible for approximately 70% of all cases (1, 2, 6). Other isolated organisms belong to the genera *Mucor*, and *Rhizomucor*; while fungi such as *Cunninghamella, Lichthemia*, and *Apophysomyces* less commonly cause infection (6). These organisms are ubiquitous in nature, found on decomposing vegetation and soil, where they grow rapidly and release large numbers of spores that can become airborne. While spores are generally harmless to immunocompetent people, almost all human infections occur in the presence of some underlying immunocompromising condition. These include hematological malignancies, organ or bone marrow transplant, corticosteroids use, hyperglycemia, diabetic ketoacidosis (DKA), and other forms of acidosis (2, 4, 8). Immunocompetent individuals suffering from burn wounds or severe trauma (e.g. soldiers in combat operations and motorcycle accident victims), or those injured in the aftermath of natural disasters (e.g., the Southeast Asian tsunami in 2004, or the tornadoes in Joplin, Missouri, in June 2011), are also uniquely susceptible to life-threatening Mucorales infections (9-11).

Devastating rhinoorbital/cerebral and pulmonary mucormycosis are the most common manifestations of the infection caused by inhalation of spores (8, 12). In healthy individuals, cilia carry spores to the pharynx, which are later cleared through the gastrointestinal tract (13). Diabetes is a risk factor that predominantly predisposes individuals to rhinoorbital/cerebral mucormycosis (RCM) (6, 8). In susceptible individuals, RCM usually begins in the paranasal sinuses, where the organisms adhere to and proliferate in the nasal epithelial cells. Eventually, adhered Mucorales invade adjoining areas such as the palate, the orbit, and the brain, causing extensive necrosis, destruction of nasal turbinates’, cranial nerve palsies and facial disfigurement, all in a short span of days to weeks. Due to the angioinvasive nature of the disease, the infection often hematogenously disseminates to infect distant organs. We have shown that *Rhizopus* thrives in high glucose, and acidic conditions and can invade human umbilical vein endothelial cells via interaction of the fungal ligand, spore-coat protein (CotH), with the host cell receptor <u>g</u>lucose <u>r</u>egulated <u>p</u>rotein <u>78</u> kDa protein (GRP78) (14, 15). In contrast, in neutropenic patients inhaled spores can directly progress into the bronchioles and alveoli causing pneumonia and rarely cause RCM (16-18). The reasons why patients with DKA are mainly infected with RCM, whereas neutropenic patients commonly suffer from pulmonary infections (8, 19) are not understood. We postulate that Mucorales ligands recognize host receptors unique to individual cell types (i.e. alveolar, nasal, endothelial cells), and that this fungal ligand-host receptor interaction is enhanced by host factors, eventually leading to infections in the respective host niches.

To investigate this hypothesis, we identified the nasal and alveolar epithelial cell receptors to Mucorales ligands and studied the effect of host factors commonly present in DKA patients on the expression and interaction of these receptors/ligands. Here we show that, similar to endothelial cells, the fungal CotH3 protein physically interacts with GRP78 on nasal epithelial cells. Elevated concentrations of glucose, iron and ketone bodies present during DKA significantly induce the expression of GRP78 and CotH3, leading to enhanced invasion and damage of nasal epithelial cells. Antibodies against either CotH3 or GRP78 abrogate *R. delemar* invasion and damage of nasal epithelial cells. In contrast, *Rhizopus* binds to integrin β1 during invasion of alveolar epithelial cells. Binding to integrin β1 triggers the activation of epidermal growth factor receptor (EGFR) signaling (20). Anti-integrin β1 antibodies significantly reduce EGFR activation, blocks alveolar epithelial cell invasion and protect neutropenic mice from pulmonary mucormycosis. These results introduce plausible explanation for the unique susceptibility of DKA patients to RCM in which inhaled Mucorales spores are trapped in the sinuses via GRP78/CotH3 overexpression. We also posit that receptors identified in this study are potential novel targets for development of pharmacologic or immunotherapeutic approaches against a variety of extremely lethal mucormycosis infections.

## Results

### Distinct host receptors are used by *R. delemar* to invade and damage nasal or alveolar epithelial cells

We compared the ability of *R. delemar* to invade and damage either nasal or alveolar A549 epithelial cells *in vitro*. Incubation of *R. delemar* germlings with either of the two cell lines, resulted in ∼40% invasion of host cells within the first 3 h of interaction, and by 6 h, almost all germlings had invaded the nasal and alveolar epithelial cells (**Fig. 1**). Interestingly, *R. delemar*-mediated damage of nasal epithelial cells occurred significantly earlier than damage of alveolar epithelial cells. Specifically, fungal germlings damaged 40% and 80% of the nasal epithelial cells within 30 h and 48 h, respectively (**Fig. 1A**). In contrast, no detectable damage and only 50% of the alveolar epithelial cells were injured after similar periods of incubation with *R. delemar* (**Fig. 1B**). These results also show that fungal invasion precedes damage of both types of epithelial cells. Importantly, *R. delemar-*mediated damage of primary human alveolar epithelial cells was similar to damage caused to A549 cells (**Fig. S1**). Therefore, the invasion and damage of the alveolar epithelial cell line is reflective of *R. delemar* interactions with primary alveolar epithelial cells.

**Fig. 1.**
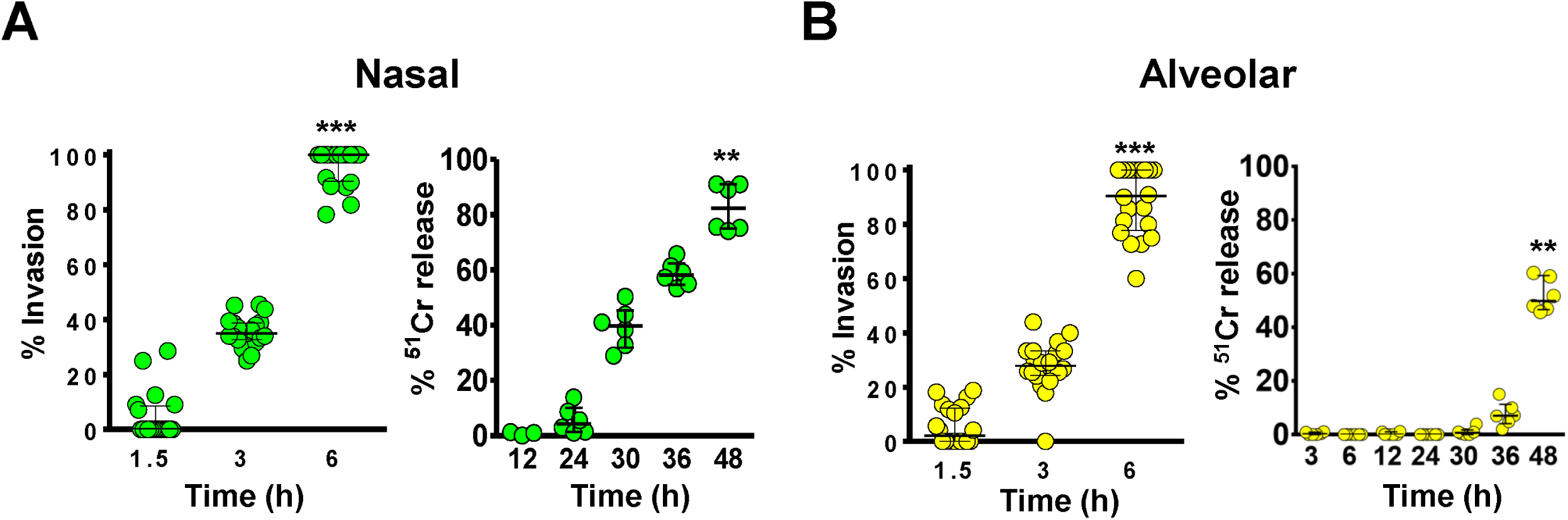
*R. delemar*-mediated invasion and damage of nasal and alveolar epithelial cells. *R. delemar* invasion of nasal (A) or alveolar (B) epithelial cells was determined using differential fluorescent assay by staining with 1% Uvitex for 1 hour, while damage assay was performed using ^51^Cr release method. *** *P* < 0.0001 and ** *P* < 0.001 compared to the first time point in each panel. Data presented as median ± interquartile range from 3 independent experiments.

We questioned if the disparity in damage to the two different epithelial cells was due to *R. delemar’s* ability to recognize different host receptors on the nasal and alveolar epithelial cells. We used an affinity purification process developed by Isberg and Leong (21), where *R. delemar* germlings were incubated separately with extracts of biotin-labeled total proteins of the nasal or alveolar epithelial cells. *R. delemar* specifically bound to a single nasal epithelial cell protein band that was isolated on an SDS-PAGE gel, and observed as a 78 kDa band post immunoblotting with anti-biotin antibodies (**Fig. 2A**). This protein band was identified by liquid chromatography–mass spectrometry (LC-MS) as the human GRP78, which we previously reported to be a receptor to invading Mucorales on human umbilical vein endothelial cells (14). To confirm the identity of the band, we stripped and probed the same immunoblot containing the nasal epithelial cell membrane proteins with an anti-GRP78 polyclonal antibodies. The polyclonal antibodies recognized the 78-kDa band that had bound to *R. delemar* germlings (**Fig. 2A**).

**Fig. 2.**
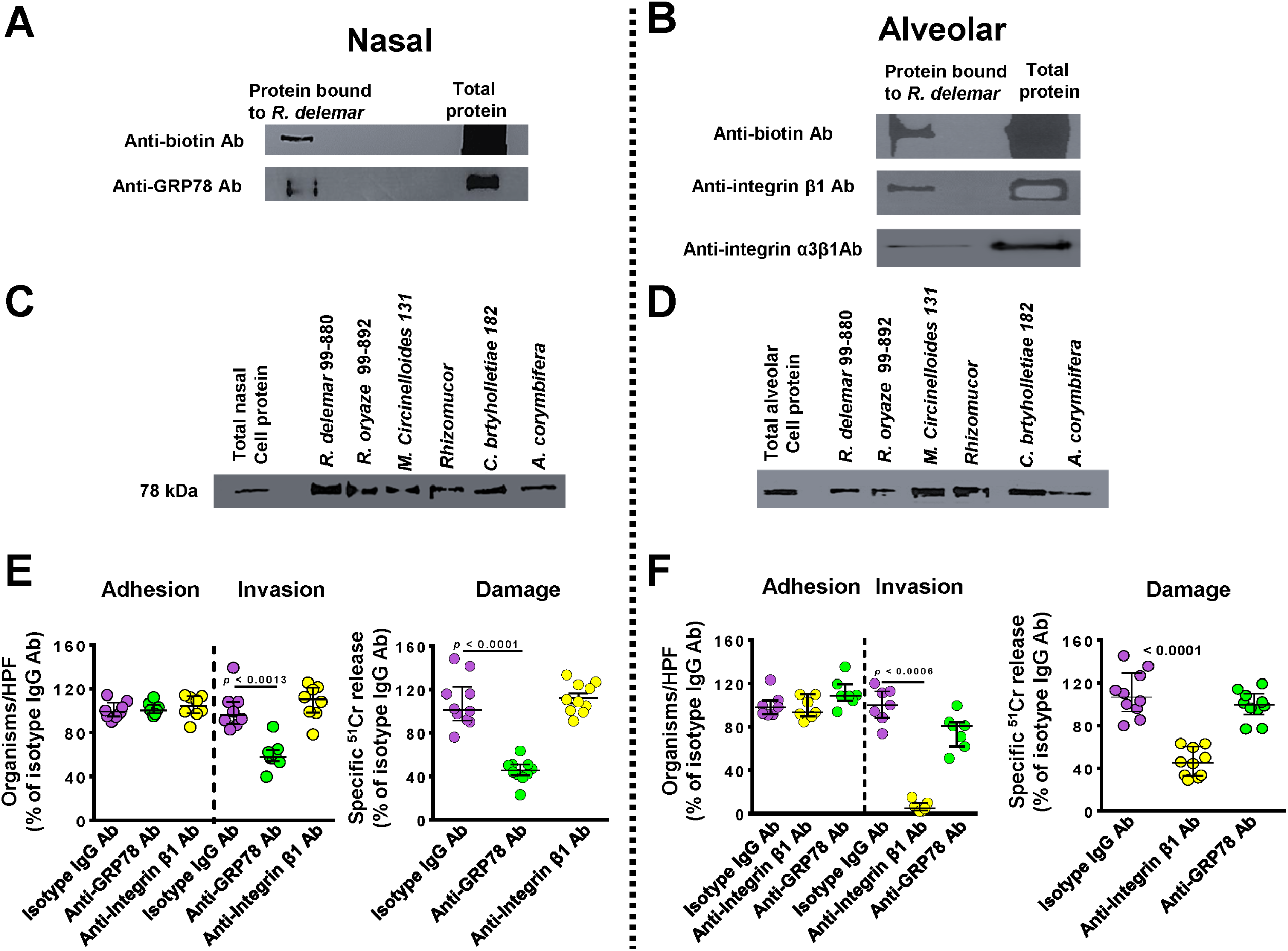
GRP78 is a nasal epithelial cell receptor, while integrin α3β1 is an alveolar epithelial cell receptor during Mucorales interaction. Biotinylated nasal (A) or alveolar (B) epithelial cells were incubated with *R. delemar* germlings and unbound proteins were removed with repeated washing. Bound proteins were separated on SDS-PAGE, and identified by Western blotting using anti-biotin monoclonal antibody (Ab, top panel) and the identity of the proteins were confirmed to be GRP78 (78 kDa) for nasal (A) or integrin β1 (130 kDa) (B) by using anti-GRP78 or anti-Integrin α3β1 antibodies, respectively (bottom panels). Affinity purification of GRP78 (C) or integrin β1 (D), respectively, by other Mucorales. Anti-GRP78 and anti-integrin antibodies block *R. delemar*-mediated invasion and subsequent damage of nasal (E) and alveolar (F) epithelial cells when compared to isotype matched-IgG, respectively. Both antibodies had no effect on adherence of the fungus to host cells. Data in (E) and (F) are expressed as median ± interquartile range from 3 independent experiments. Different color codes are used to simplify the graph; purple, isotype IgG; green, anti-GRP78 Ab; and yellow, anti-integrin β1 Ab.

Similarly, only a single 130 kDa protein band from the alveolar epithelial cell extracts was bound to *R. delemar* germlings (**Fig. 2B**). This protein was identified as integrin β1 by LC-MS, and subsequently confirmed by probing with an anti-integrin β1 antibody on Western blotting (**Fig. 2B**). Integrins are known to be highly expressed in human lung tissues (https://www.ncbi.nlm.nih.gov/gene/3675), and we found that gene expression of integrin β1, but not GRP78, was upregulated in alveolar epithelial cells during infection with *R. delemar* (**Fig. S2**). Furthermore, transcriptomic analysis of mouse lung tissues in early stages of pulmonary mucormycosis identified an upregulation of a gene encoding for integrin α3 (22). Since integrins function as heterodimers (23), we sought to verify if integrin α3 subunit combines with integrin β1 in acting as a putative receptor to *R. delemar* by alveolar epithelial cells. An integrin α3β1 polyclonal antibody recognized the 130 kDa band from A549 alveolar epithelial cells (**Fig. 2B**). Therefore, it is possible that α3 subunit functions as a heterodimer with β1 to serve as a receptor during Mucorales invasion of alveolar epithelial cells.

To investigate if nasal GRP78 and alveolar integrin α3β1 are putative universal receptors to other Mucorales, we performed the affinity purification experiment using germlings of other Mucorales clinical isolates. Indeed, all tested Mucorales including *R. oryzae* 99-892, *M. circinelloides* 131, *Rhizomucor, C. bertholletiae* 182 and *L. corymbifera* 008-0490 bound GRP78 and integrin α3β1 from nasal (**Fig. 2C**) and alveolar (**Fig. 2D**) epithelial cells, respectively. Collectively, these data suggest that Mucorales interacts with nasal and alveolar epithelial cells by using different host receptors.

### GRP78 and integrin β1 are receptors on nasal and alveolar epithelial cells, respectively

To confirm the function of GRP78 and integrin β1 on nasal and alveolar epithelial cells as receptors to *R. delemar*, we examined the effect of anti-GRP78 and anti-integrin β1 antibodies on *R. delemar-*mediated host cell adhesion, invasion and subsequent damage. While incubating nasal epithelial cells with anti-GRP78 polyclonal antibodies resulted in ∼50% inhibition of *R. delemar*-mediated host cell invasion, the antibodies had no effect on adhesion when compared to isotype-matched control antibodies (**Fig. 2E**). The anti-GRP78 antibodies also reduced *R. delemar* ability to injure nasal epithelial cells by ∼60%. As expected, anti-integrin β1 antibodies had no effect on *R. delemar–*mediated adhesion to, invasion and damage of nasal epithelial cells (**Fig. 2E**). In contrast, when compared to isotype-matched control antibodies, the use of anti-integrin β1 antibodies, and not anti-GRP78 antibodies, almost completely abolished the ability of *R. delemar* to invade alveolar epithelial cells (>95% reduction in invasion) (**Fig 2F**). Similar to anti-GRP78 and nasal epithelial cells, anti-integrin β1 antibodies had no effect on the adherence of the fungus to alveolar epithelial cells (median adherence of 98%, 93% and 108% for isotype-matched IgG, anti-GRP78 IgG, anti-integrin β1 IgG, respectively, *P*>0.1). Finally, only anti-integrin β1 antibodies decreased the mold-mediated damage to alveolar epithelial cells by ∼60% (**Fig. 2F**). Overall, these results highlight that GRP78 and integrin β1 act as major and specific receptors to *R. delemar* during invasion and subsequent damage of nasal and alveolar epithelial cells, respectively.

We previously demonstrated the importance of *R. delemar* interacting with GRP78 by overexpressing GRP78 on Chinese Hamster Ovarian cells (CHO) and showed increased *R. delemar-*mediated invasion and damage of the transfected cells (14). To validate the importance of integrin β1 as a receptor for *R. delemar* during invasion of alveolar epithelial cells, we compared the ability of *R. delemar* to invade and damage GD25 fibroblast cell line which is generated from an integrin β1^-/-^ mouse, to β1AGD25 cell line made by transfecting GD25 cells with mouse integrin β1 cDNA (24). Despite the lack of difference in adhesion of *R. delemar* to these two cell lines, the β1GD25 fibroblast cells expressing integrin β1 was more susceptible to *R. delemar*– mediated invasion and damage when compared to GD25 cells lacking integrin β1 (an increase of ∼ 600% for invasion and 150% for damage of β1GD25 versus GD25 cells) (**Fig. 3A**). These data reaffirm the importance of integrin β1 as a host receptor for *R. delemar* during invasion and subsequent damage of alveolar epithelial cells.

**Fig. 3.**
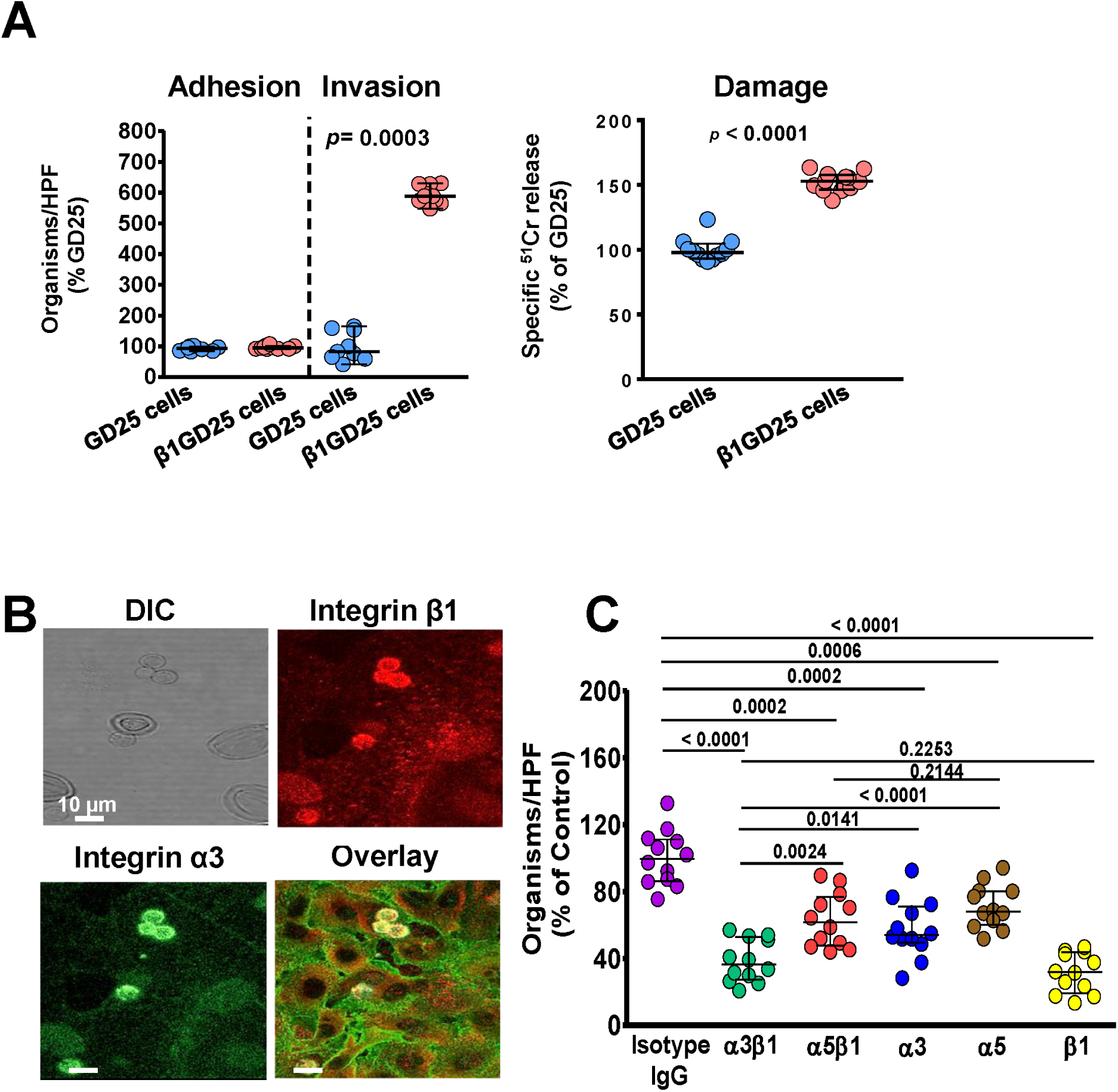
Integrin α3β1 is required for *R. delemar*-mediated host cell invasion and damage. *R. delemar has* reduced invasion and damage of GD25 fibroblast cell line lacking integrin β1, compared to β1GD25, an integrin β1 restored fibroblast cell line (A). Adhesion and invasion of GD25 and β1AGD25 fibroblast cell lines were conducted using differential fluorescent assay, while host cell damage was carried out using ^51^Cr release method. Confocal microscopy images showing the accumulation of integrin α3β1 around *R. delemar* during infection of alveolar epithelial cells (B). Images were taken after 2.5 hours of incubation of the fungus with the host cells. Anti-integrin α3β1 monoclonal antibody block *R. delemar*-mediated invasion of alveolar epithelial cells (C). Alveolar epithelial cells were incubated with 5 µg/ml of different anti-integrins antibodies or isotype-matched IgG for 1 hour prior to infecting with *R. delemar.* Data are expressed as median ± interquartile range from 3 independent experiments for (A) and (C).

For cell membrane proteins to act as host cell receptors they must be in close proximity to invading fungal cells. Therefore, we used an indirect immunofluorescence assay to localize integrin α3β1 on alveolar epithelial cells during infection with *R. delemar* germlings. Both integrin α3 (stained with anti-intgerin α3 antibody fluorescing green) and β1 (stained with anti-integrin β1 antibody fluorescing red) were expressed on the surface of alveolar epithelial cells and coalesced on invading *R. delemar* germlings with an overlay images showing clear intense yellow staining around the fungal cells (**Fig. 3B**).

We previously showed that the filamentous fungal pathogen *A. fumigatus* invades alveolar epithelial cells through the fungus CalA protein binding to integrin α5β1 (25). Thus, to evaluate the function of integrin α5 as a potential receptor for *R. delelmar*, we repeated the indirect immunofluorescence assay using antibodies targeting integrin β1 and α5. As expected, integrin β1 accumulated as a distinct ring-like formation around endocytosed *R. delemar* germlings. In contrast, integrin α5 had a diffused staining without accumulation around invading germlings (**Fig. S3**). Thus, these data strongly suggest that the receptor for *R. delemar* during invasion of alveolar epithelial cells is likely to be integrin α3β1, rather than α5β1.

To confirm the identity of the alveolar epithelial cell receptor during Mucorales invasion, we incubated the *R. delemar* germlings with A549 epithelial cells in the presence of specific monoclonal antibodies targeting either integrin α3, α5, or β1 separately and the two dimers of integrin α3 β1, or α5 β1. While all treatments resulted in reduction of cellular invasion compared to the isotype-matched IgG antibodies (which did not block invasion), there were differences in the extent of invasion inhibition. Specifically, targeting integrin β1 caused the greatest reduction in invasion with ∼70% inhibition, while anti-integrin α3 and anti-integrin α5 antibodies individually provided ∼50% and 30% protection from invasion, respectively (**Fig. 3C**). Interestingly, targeting both integrin α3β1 resulted in similar inhibition of *R. delemar* invasion of A549 cells when compared to invasion inhibition provided by anti-β1 antibody of ∼70% and significantly more than the invasion inhibition generated by anti-α3 antibody or anti-α5β1 (**Fig. 3C**). Collectively, these results show that integrin β1 is the major host receptor acting as a heterodimer with α3 during *R. delemar* invasion of alveolar epithelial cells and blocking these receptors can reduce *R. delemar* virulence to alveolar epithelial cells *in vitro*.

### Integrin β1 signaling is required for EGFR phosphorylation in alveolar epithelial cells during Mucorales infection

We recently reported EGFR acts as a receptor for *R. delemar* during invasion of alveolar epithelial cells (20). However, the mechanism by which EGFR signaling is stimulated during infection was not identified. We tested if integrin β1 signaling played a role in stimulating EGFR activation during *R. delemar-*invasion, by examining phosphorylation of the A549 cells’ EGFR tyrosine residue 1068 in the presence of anti-integrin β1 antibodies. Using immunoblotting assay, we determined that infection with *R. delemar* induces the EGFR phosphorylation in A549 cells. When the *R. delemar*/A549 cell interaction was performed in the presence of integrin β1 antibodies, the phosphorylation of EGFR was abolished to the basal levels (**Fig. 4**). Thus, these results are consistent with a model in which *R. delemar* interacts with integrin β1 causing activation of EGFR.

**Fig. 4.**
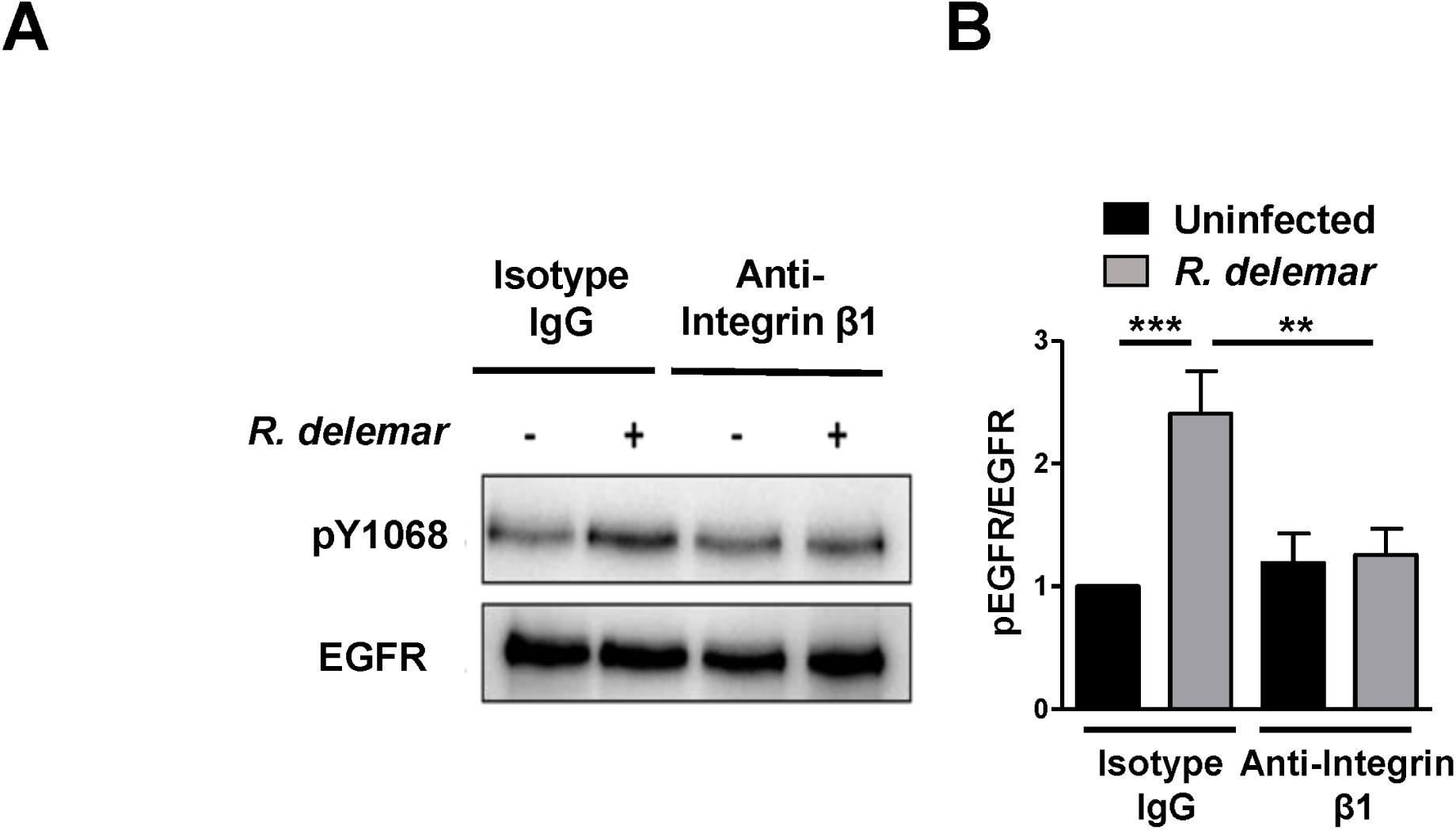
Anti-integrin antibodies block activation of alveolar epithelial cell EGFR. Representative immunoblots (A) and densitometric analysis (B) show that *R. delemar* infection induced phosphorylation of EGFR on tyrosine residue 1068 compared to control and anti-integrin β1 antibody blocked it. Data in (B) are mean ± SD of three independent experiments.

### *R. delemar* cell surface proteins CotH3 and CotH7 are the fungal ligands to nasal and alveolar epithelial cells, respectively

Having identified the receptor on nasal and alveolar epithelial cells that interacts with *R. delemar* germlings, we next sought to identify the fungal cell surface protein that binds to GRP78 and integrin α3β1. Far-Western blot analysis using recombinant human GRP78 followed by anti-GRP78 antibodies or human integrin α3β1 followed by anti-integrin α3β1 antibodies identified the presence of prominent bands from the supernatant of *R. delemar* regenerated protoplasts that bound to GRP78 (**Fig. 5A**) or integrin α3β1 (**Fig. 6A**). LC-MS of the bands identified CotH3 and CotH7 as putative fungal ligands binding to GRP78 and integrin α3β1, respectively. We previously described that CotH3 is the fungal ligand to host GRP78 during interaction of *R. delemar* with human umbilical vein endothelial cells (15, 26). Therefore, we used tools available for us to determine the importance of CotH3 to *R. delemar* when interacting with nasal epithelial cells. We incubated biotinylated nasal epithelial cell membrane proteins with the model yeast *Saccharomyces cerevisiae* harboring a plasmid expressing CotH3 or *S. cerevisiae* expressing the empty plasmid as a negative control. The CotH3 expressing *S. cerevisiae* bound the 78-kDa protein of GRP78 as confirmed by Western blotting with anti-GRP78 antibodies, whereas *S. cerevisiae* strain expressing empty plasmid did not (**Fig. 5B**). Next, we visualized the interaction between the two host-fungal proteins by a proximity ligation assay (PLA). In this assay, non-fluorescent primary antibodies (commercially available) raised in different species are allowed to recognize GRP78 and CotH3 (using anti-CotH3 antibodies that we previously described (27)) on the host cells and fungus, respectively. Secondary antibodies directed against the constant regions of the two primary antibodies called PLA probes bind to the primary antibodies. The PLA probes fluoresce as a distinct bright spot only if the two proteins of GRP78 and CotH3 are in close proximity. Indeed, nasal epithelial cell-*R. delemar* germling interaction triggered the probe to fluoresce red (**Fig. 5C**). This fluorescence was located on germlings that interacted with host cells stained with DAPI yielding a bright pink color. Therefore, *R. delemar* CotH3 interacts with the GRP78 receptor on nasal epithelial cells leading to invasion and subsequent damage of host cells.

**Fig. 5.**
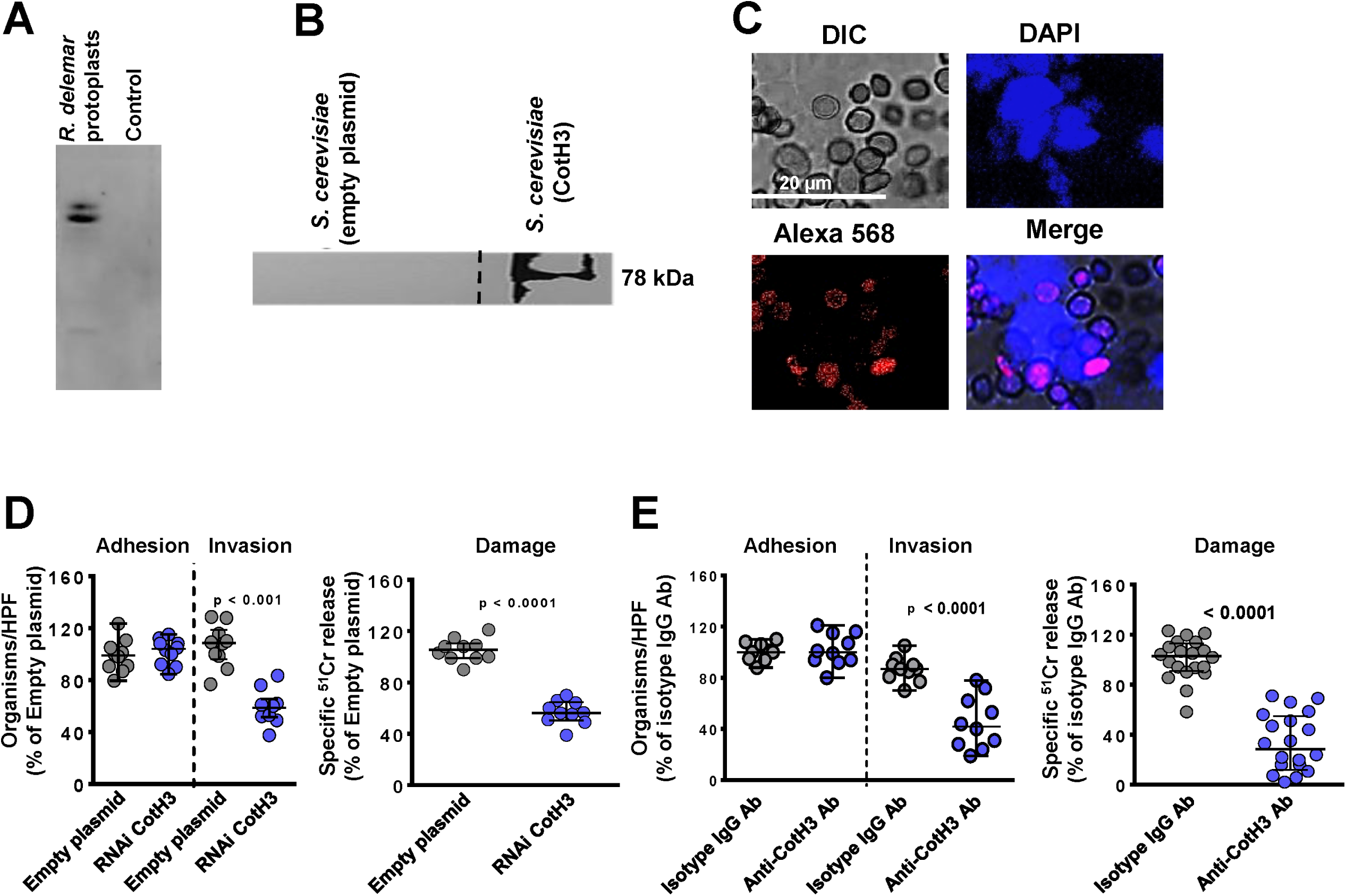
CotH3 is the *R. delemar* cell-surface ligand to GRP78 on nasal epithelial cells. Far-Western blot of *R. delemar* surface proteins that bound to GRP78 (A). Affinity purification of nasal cell GRP78 by *S. cerevisiae* cells expressing CotH3 identified by anti-GRP78 antibody (B). Dashed line represent cropped image from Fig. 2A GRP78 blot. Confocal microscopy images showing interaction of nasal epithelial GRP78 and *R. delemar* CotH3 after 2.5 hours incubation shown by proximity ligation assay (PLA) (C). DAPI staining was used to identify host cells. Inhibition of CotH3 expression by RNAi reduced the ability of *R. delemar* to invade (by differential fluorescence) and damage (by ^51^Cr release method) nasal epithelial cells compared with empty plasmid transformed *R. delemar* (D). Anti-CotH3 antibody blocked *R. delemar-*mediated invasion of and damage to nasal epithelial cells. Data in (D) and (E) are expressed as median ± interquartile range from 3 independent experiments.

**Fig. 6.**
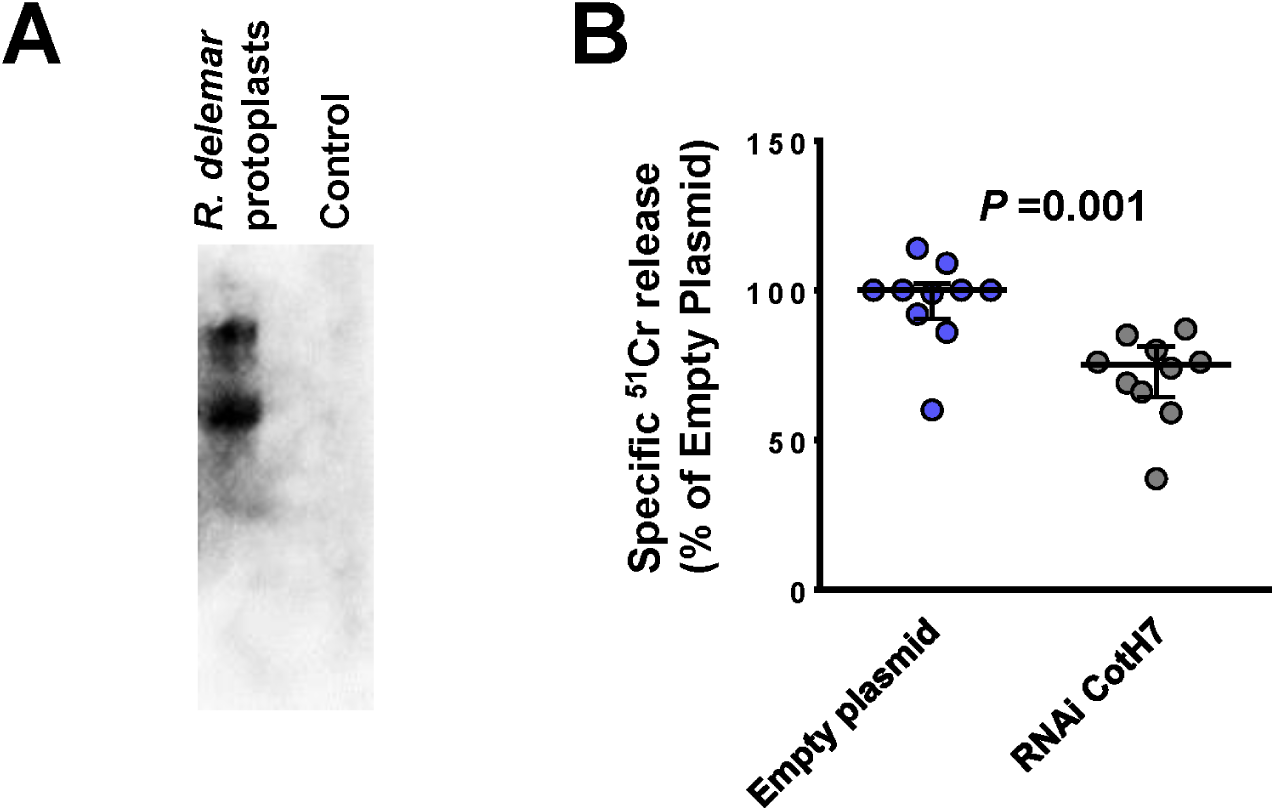
CotH7 is the *R. delemar* cell-surface ligand to integrin α3β1. Far-Western blot of *R. delemar* surface proteins that bound to integrin (A). Inhibition of CotH3 expression by RNAi reduced the ability of *R. delemar* to invade (by differential fluorescence) and damage (by ^51^Cr release) alveolar epithelial cells compared with empty plasmid transformed *R. delemar* (B). Data in (B) are expressed as median ± interquartile range from 3 independent experiments.

To investigate if the interactions of CotH3/GRP78 and CotH7/integrin α3β1 result in mediating *R. delemar* invasion and damage of nasal and alveolar epithelial cells, we specifically down regulated the expression of CotH3 or CotH7 in *R. delemar* by RNAi. Individually targeting CotH3 and CotH7 by RNAi resulted in generating *R. delemar* mutants that had ∼90% (15) and 50% (**Fig. S4**) inhibition in these two genes, respectively. Mutants were compared to the *R. delemar* strain transformed with an empty plasmid, in their ability to invade and damage nasal and alveolar epithelial cells. Incubating nasal epithelial cells with an *R. delemar* RNAi-suppressed CotH3 strain displayed >50% defect in invasion of and damage to nasal epithelial cells when compared to *R. delemar* strain transformed with the empty plasmid. The inhibition of CotH3 expression had no effect on adherence of *R. delemar* to nasal epithelial cells (**Fig. 5D**), nor did it affect the ability of *R. delemar* to interact with alveolar epithelial cells (**Fig. S5**). Therefore, CotH3 is a specific *R. delemar* ligand that mediates invasion and subsequent damage to nasal epithelial cells..

We previously generated anti-CotH3 antibodies that blocked *R. delemar* mediated invasion of endothelial cells. Therefore, we tested the ability of these antibodies to block *R. delemar-*mediated invasion of and subsequent damage to nasal epithelial cells. Anti-CotH3 antibodies resulted in 60% and 75% reduction in the ability of *R. delemar* to invade and damage nasal epithelial cells when compared to isotype-matched control IgG, respectively (**Fig. 5E**). These results further confirm the importance of CotH3 protein in *R. delemar* interacting with nasal epithelial cells *in vitro.* Interestingly, anti-CotH3 antibodies showed reduction in *R. delemar*-mediated invasion to alveolar cells when compared to isotype-matched IgG (**Fig. S5B**).

Down regulation of CotH7 expression resulted in a statistically significant reduction (30% reduction) in *R. delemar-*mediated damage of alveolar epithelial cells (**Fig. 6B**). Similar to the outcome of RNAi CotH3 mutant interacting with alveolar epithelial cells, down regulation of the CotH7 expression had no effect on *R. delemar* interacting with nasal epithelial cells (**Fig. S7**). Therefore, interactions of *R. delemar* with alveolar epithelial cells are mainly driven by CotH7 binding to integrin α3β1.

### DKA host factors enhance *R. delemar-*mediate damage of nasal but not alveolar epithelial cells

We have previously shown that endothelial cell GRP78 and Mucorales CotH3 are overexpressed in physiological conditions found in DKA patients such as hyperglycemia, elevated available serum iron and high concentrations of ketone bodies, leading to enhanced invasion and damage of endothelial cells (14, 15, 26). Because we found that *R. delemar* uses similar mechanism to interact with nasal epithelial cells, we reasoned that upregulation of GRP78 on nasal epithelial cells might lead to entrapment of inhaled spores in the nasal cavity of DKA patients leading to rhino-orbital disease rather than pulmonary infection. To test this hypothesis, we measured the effect of physiologically elevated concentrations of glucose, iron and β-hydroxy butyrate (BHB, as a representation for ketone bodies) on the GRP78 expression of nasal epithelial cells and subsequent interactions with *R. delemar.* The use of elevated concentrations of glucose (4 or 8 mg/ml), iron (15-50 μM of FeCl_3_), or BHB (5-10 mM) resulted in ∼2-6 fold increase in the surface expression of GRP78 on nasal epithelial cells when compared to normal concentrations of 1 mg/ml glucose, 0 μM iron, or 0 mM BHB (**Fig. 7A**). This enhanced expression of GRP78 coincided with increased ability of *R. delemar* to invade (**Fig. 7B**) and subsequently damage (**Fig. 7C**) nasal epithelial cells (∼150%-170% increase in invasion and 120%-170% in nasal epithelial cells damage vs. normal concentration of the effector). Conversely, the same elevated concentrations of glucose, iron and BHB had no effect on the surface expression of integrin β1 of alveolar epithelial cells (**Fig. 8A**) nor did it result in enhanced *R. delemar-*mediated invasion (with the exception of 8 mg/ml glucose that caused a modest increase in invasion of 25% versus 1 mg/ml glucose) (**Fig. 8B**). Surprisingly, and in general, elevated concentrations of glucose, iron, or BHB resulted in 40-50% protection of alveolar epithelial cells from *R. delemar*-mediated injury (**Fig. 8C**). Collectively, these data suggest that nasal epithelial cells are more prone to *R. delemar-*-mediated invasion and injury than alveolar epithelial cells when exposed to DKA host factors, and likely explain at least, in part, the reason why DKA patients predominantly suffer from rhinoorbital rather than pulmonary mucormycosis.

**Fig. 7.**
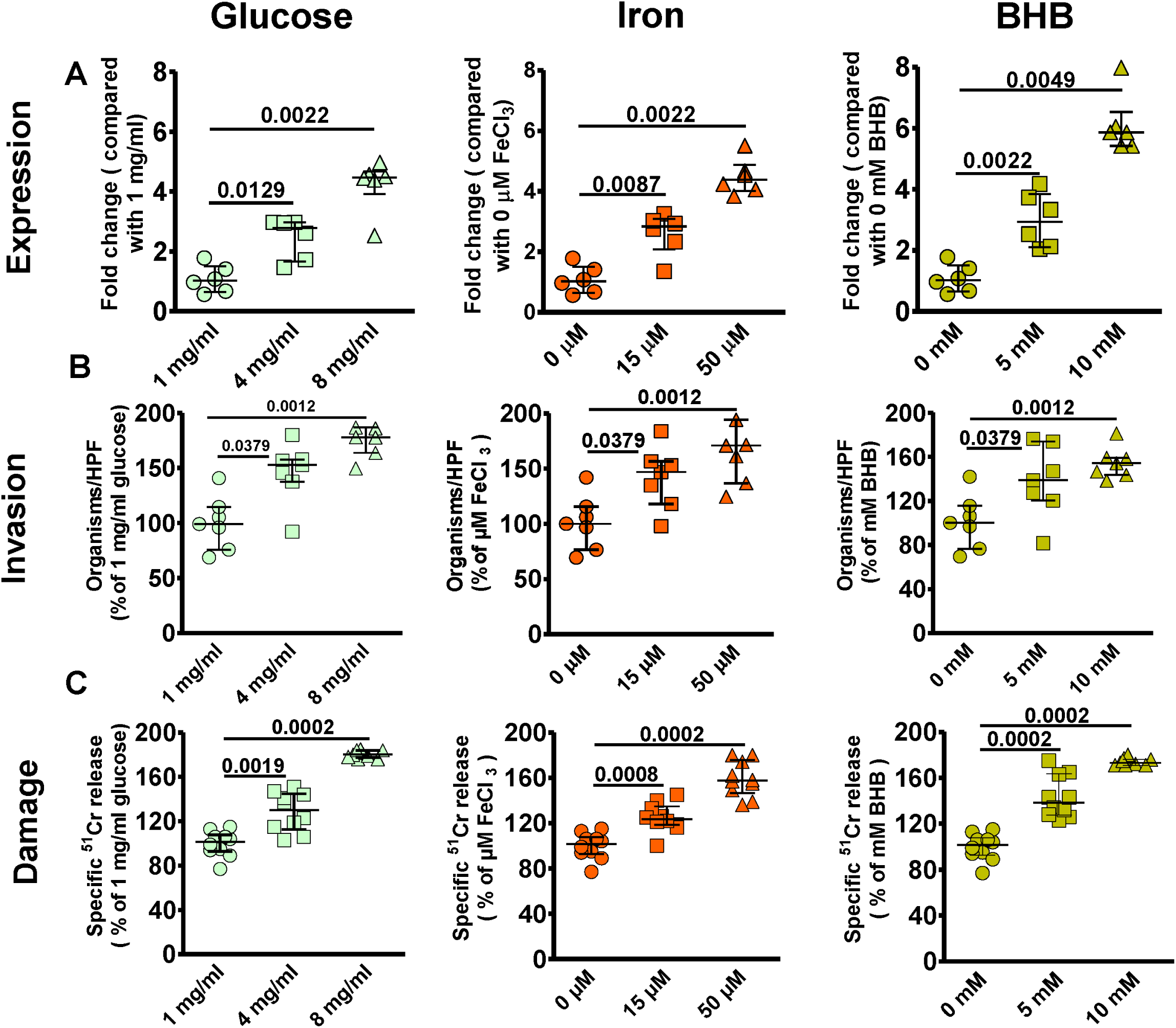
DKA host factors increase nasal epithelial cell GRP78 expression and host cell susceptibility to *R. delemar*–mediated invasion and damage. Nasal epithelial cells were incubated with physiologically elevated concentrations of glucose, iron or BHB for 5 hours and GRP78 gene expression determined by qRT-PCR (A). Elevated concentrations of glucose, iron or BHB significantly enhanced *R. delemar*-mediated nasal epithelial cell invasion (B) and damage (C). Fold changes were calculated by comparison to the lowest concentration of the exogenous factors used. Data are expressed as median ± interquartile range from 3 independent experiments.

**Fig. 8.**
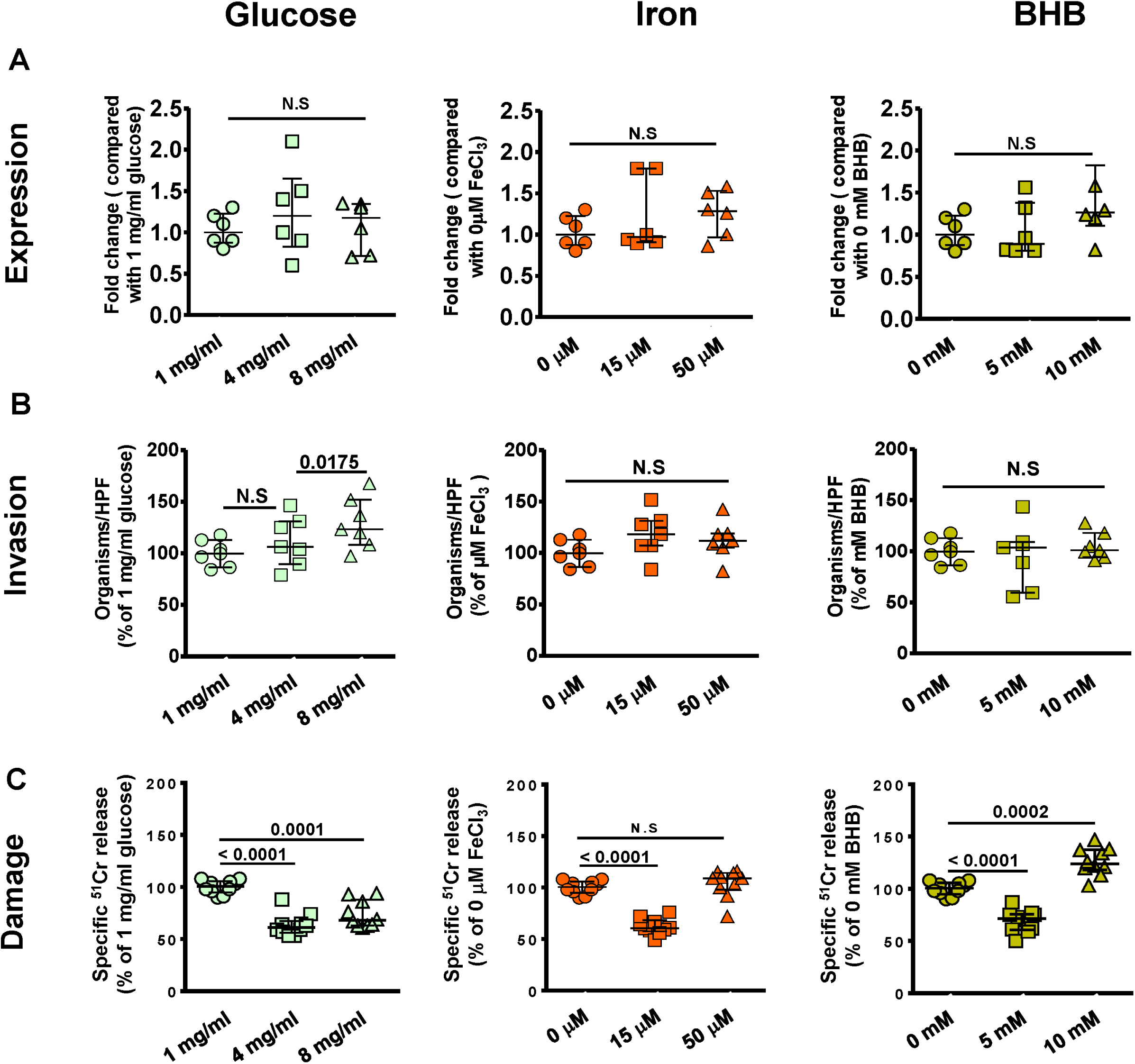
DKA host factors have no effect on integrin β1 expression levels nor they affected *R. delemar* interactions with alveolar epithelial cells. Alveolar epithelial cells were incubated with physiologically elevated concentrations of glucose, iron or BHB for 5 hours and integrin β1 gene expression determined by qRT-PCR (A). Elevated concentrations of glucose, iron or BHB had no effect on *R. delemar*–mediated alveolar epithelial cell invasion (B) or subsequent damage (C). Data are expressed as median ± interquartile range from 3 independent experiments.

### Anti-Integrin β1 antibodies protect neutropenic mice from pulmonary mucormycosis

We have previously shown that GRP78 can be targeted for treating experimental mucormycosis (14). To examine the potential of targeting integrins in treating pulmonary mucormycosis, we infected neutropenic mice intratracheally with *R. delemar* spores, and treated them one day after infection with either an isotype-matched IgG or anti-integrin β1 polyclonal IgG. While mice treated with the isotype-matched IgG had a median survival time of 11 days and 100% mortality by day 15 post infection, mice treated with the anti-integrin β1 IgG had an improved median survival time of 16 days and 30% of the mice survived by day 21 post infection when the experiment was terminated (**Fig. 9**). The surviving mice appeared healthy, and lungs and brains (primary and secondary target organs in this model (28)) harvested from the surviving mice had no residual infection as determined by lack of fungal growth from harvest organs when cultured on potato dextrose agar (PDA) plates. Thus, these data suggest that targeting integrin β1 should be explored to serve as a promising novel therapeutic option against pulmonary mucormycosis.

**Fig. 9.**
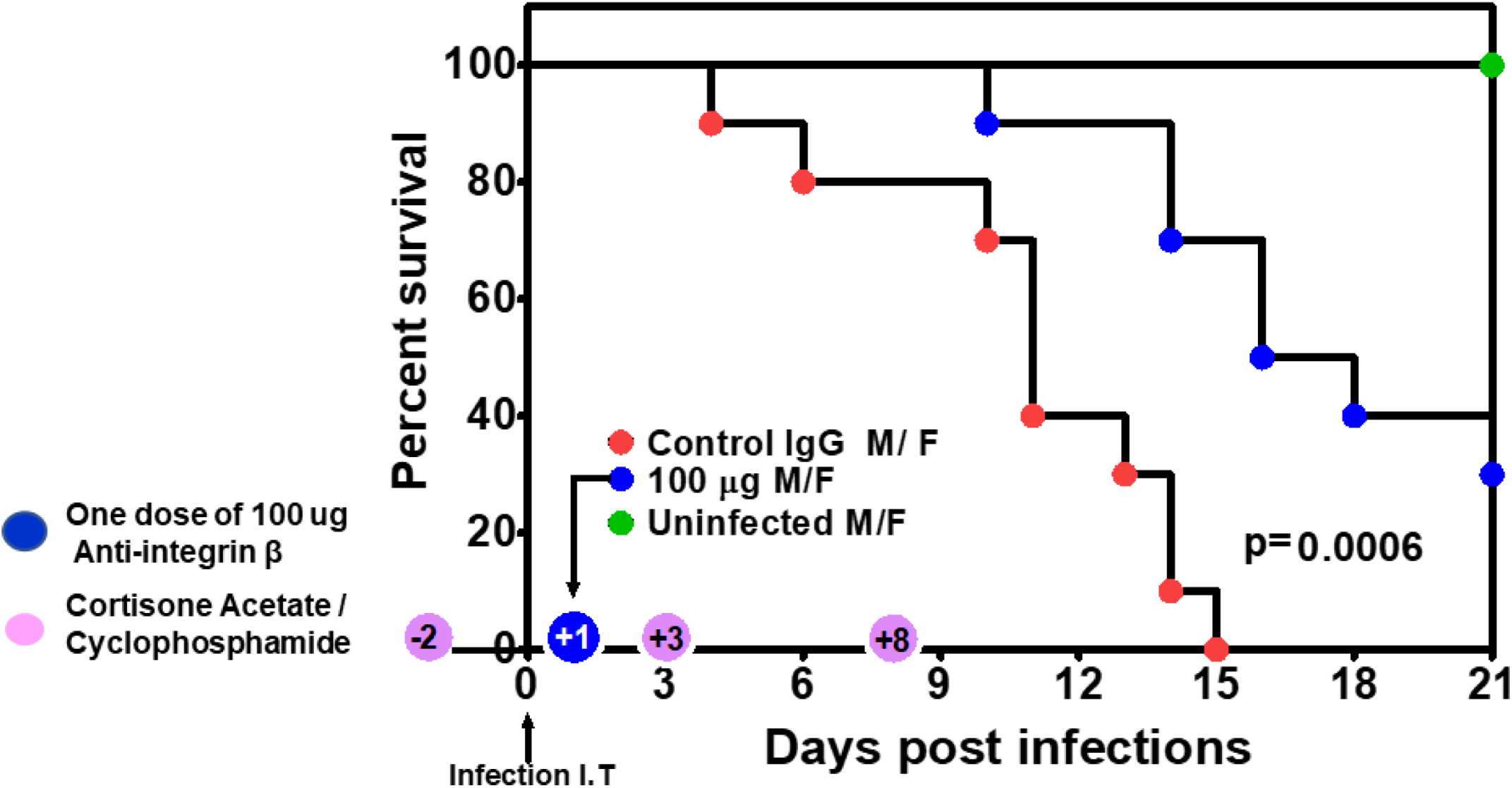
Anti-integrin β1 antibodies protect immunosuppressed mice from invasive pulmonary mucormycosis due to *R. delemar.* ICR mice (n= 10 [5 female and 5 male]/group with no difference in survival among the two genders]) were immunosuppressed on day -2, +3 and +8 with cyclophosphamide and cortisone acetate and infected on day 0 intratracheally with *R. delemar* (actual inhaled inoculum of 2.8 × 10^3^/mouse). Twenty four hours post infection, mice were treated with a single dose of either a 100 µg of an isotype-matched IgG (Control) or an anti-integrin β1 antibody. *P* = 0.0006 by Log-rank test.

## Discussion

Rhinoorbital/cerebral and pulmonary infections are the two most common manifestations of lethal mucormycosis (29). Despite acquiring the infection through inhaled spores, these two forms of disease manifestations are determined by host underlying predisposing factors. Specifically, patients with DKA appear to be more likely than other susceptible hosts to have rhinoorbital/cerebral infection, while pulmonary mucormycosis afflicts neutropenic/leukemic hosts (12, 30, 31). Since the reason for this disparity is unknown (32), we questioned if Mucorales recognize host receptors expressed uniquely in distinct niches, especially in response to specific host environmental conditions. We previously found that the fungal cell surface CotH3 protein, a unique invasin to Mucorales fungi, binds to mammalian GRP78 when infecting and damaging umbilical vein endothelial cells (14, 15). Importantly, the expression of GRP78 host receptor and CotH3 fungal ligand increases several folds under physiological conditions present in the DKA patients such as hyperglycemia, elevated iron, and ketoacidosis leading to enhanced fungal invasion, subsequent damage of endothelial cells and disease progression (26). Similar to these findings, we present multiple evidences by using affinity purification, specific antibody blocking, colocalization and gene downregulation studies, to show that *R. delemar* invades and damages nasal epithelial cells by CotH3 interacting with GRP78. As expected, DKA conditions of hyperglycemia, elevated iron, and ketoacidosis resulted in upregulation of GRP78 of nasal epithelial cells causing enhanced fungal invasion. Therefore, in patients with DKA, inhaled Mucorales spores are likely trapped in the nasal milieu by the interaction of upregulated expression of GRP78/CotH3 resulting in rhinoorobital mucormycosis (**Fig. 10A**). The highly angioinvasive *R. delemar* can eventually spread from the damaged nasal epithelial cells into surrounding tissue vasculature by continuing to interact with GRP78 on endothelial cells (15, 26). In contrast, by using similar approaches we show that the integrin α3β1 is the receptor for *R. delemar* on alveolar epithelial cells which activated EGFR resulting in invasion and pulmonary infection (**Fig. 10B**). However, hyperglycemia, elevated iron, and ketoacidosis, as seen in DKA patients did not increase integrin α3β1 expression on alveolar epithelial cells. In fact, through an unexplained mechanism(s), elevated physiological concentrations of glucose, iron and BHB protected A549 cells from invasion and subsequent damage by *R. delemar*. The protection of alveolar epithelial cells from *R. delemar-*mediated invasion and subsequent damage when exposed to elevated glucose, iron, or BHB, likely to provide further explanation on why DKA patients rarely develop pulmonary disease. Future studies will investigate the mechanism by which DKA host factors protect alveolar epithelial cells from *R. delemar* –mediated invasion and damage.

**Fig. 10.**
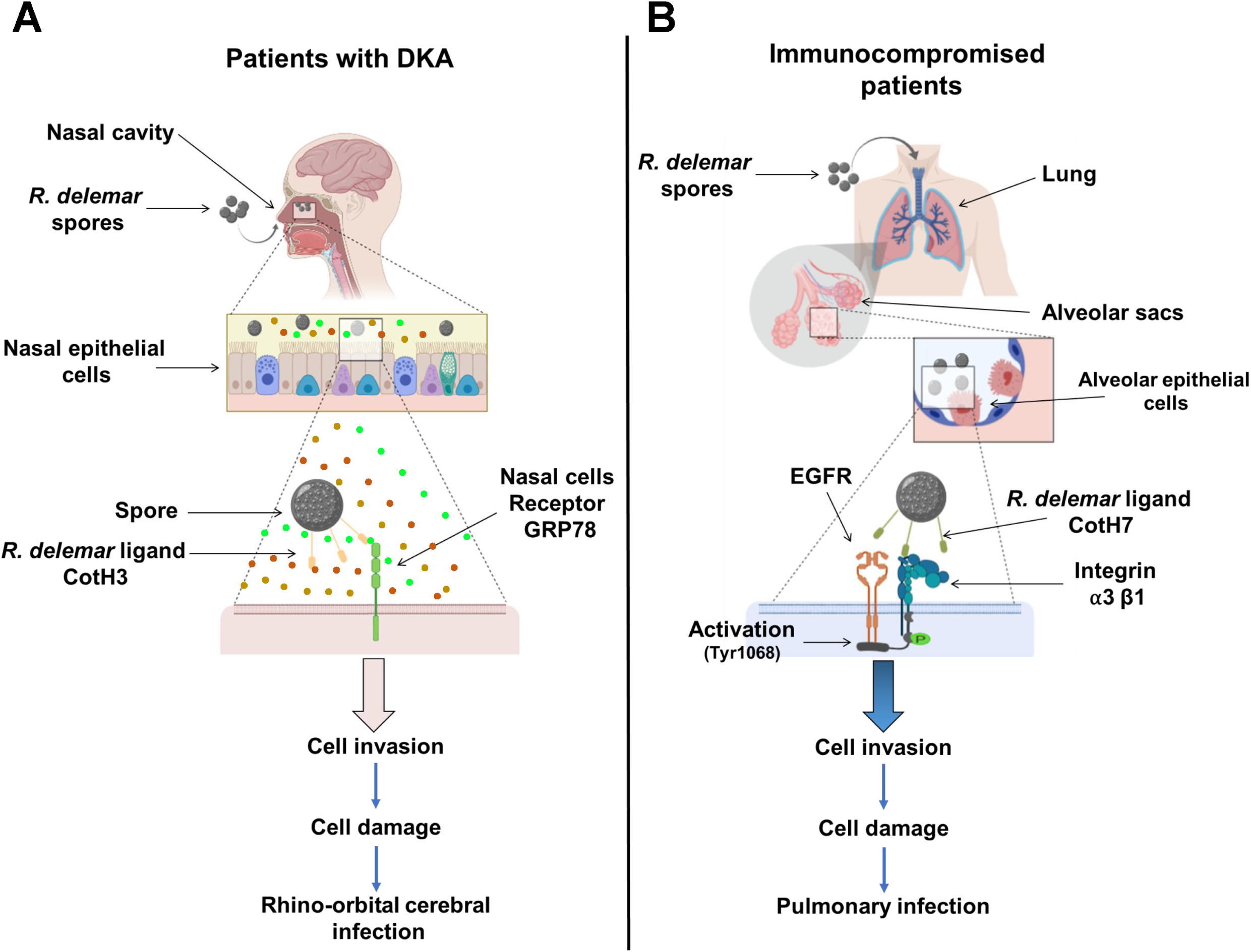
A diagram showing the molecular pathogenesis of the two main manifestation of mucormycosis. *R. delemar* inhaled spores are trapped in the sinus cavity of patients with DKA due to the overexpression of GRP78 on nasal epithelial cells and the interaction with fungal CotH3 resulting in rhinoorbital/cerebral mucormycosis (A). In immunosupporesed patients, inhaled spores reach the alveoli and bind to integrin α3β1 via fungal CotH7, thereby triggering activation of EGFR and subsequent invasion and pulmonary infection (B).

One of the intriguing results is the difference of susceptibility of nasal and alveolar epithelial cells to *R. delemar*-mediated damage despite equally being susceptible to fungal invasion. Specifically, nasal epithelial cells were more susceptible to fungal damage when compared to alveolar epithelial cells (**Fig. 1**). We previously reported on the role of *R. delemar* toxins in mediating damage to host cells (33). It is possible that the two cell types have distinct susceptibility and/or induce different levels and/or types of these toxins. Alternatively, binding to distinct receptors is likely to induce specific signal transductions pathways that might explain the differences in host cell death pattern. These possibilities are the topic of active investigation in our laboratory.

We previously reported on the CotH gene family which is uniquely and universally present in Mucorales fungi and required for mucormycosis pathogenesis (15, 22). Specifically, CotH3 mediates invasion of endothelial cells by binding to GRP78 (15, 26). *R. delemar* also uses CotH3 to invade nasal epithelial cells via binding to GRP78. However, in lung tissues where integrins are highly expressed (https://www.ncbi.nlm.nih.gov/gene/3675), CotH7 appears to be the major *R. delemar* ligand mediating binding to integrin α3β1 of alveolar epithelial cells. Although CotH2 and CotH3 proteins are closely related, CotH7 is distantly related with ∼ 50% amino acid identity to CotH3 (**Fig. S7**). It is noted that CotH2, CotH3, and CotH7 are among the most expressed genes in the entire genome of two clinical isolates (*R. delemar* 99-880 and *R. oryzae* 99-892) and their expression is not induced by alveolar epithelial cells (22). This non-induced high expression and the presence of altered protein family members is likely necessary for the organism to successfully infect host niches in which invasion of tissues is dictated by the presence of different receptors. However, in both nasal and alveolar epithelial cells, antibody blocking studies targeting the receptors or the ligands did not completely block *R. delemar-*mediated adhesion, invasion or damage of host cells. Thus, other host receptors/fungal ligands are likely to be involved in these interactions.

We found that anti-CotH3 antibodies, but not reduction of CotH3 expression by RNAi, were able to block invasion, and to a lesser extent, adherence of *R. delemar* to alveolar epithelial cells (**Fig S5**). It is noted, that the antibodies were generated against a peptide of CotH3 (MGQTNDGAYRDPTDNN (27)) that is ∼70% conserved in CotH7 protein (Fig S8), whereas the inhibition of CotH3 expression by RNAi resulted in ∼ 80% gene silencing (15).

Integrins are a family of adhesion receptors consisting of α and β heterodimer transmembrane subunits that are specialized in binding cells to the extracellular matrix (23). They can also function as receptors for extracellular ligands and transduce bidirectional signals into and outside the cell using effector proteins (34, 35). One of such pathways is the ability of integrins to cooperate with EGFR leading to synergy in cell proliferation, cell survival, and cell migration (36). We recently reported on the use of an unbiased survey of the host transcriptional response during early stages of *R. delemar* infection in a murine model of pulmonary mucormycosis as well as an *in vitro* A549 cell infection model by using transcriptome analysis sequencing. RNA-seq data showed an activation of the host’s EGFR by an unknown mechanism (20). Furthermore, an FDA-approved inhibitor of EGFR, gefitinib, successfully inhibited alveolar epithelial cell invasion by *R. delemar in vitro* and ameliorate experimental murine pulmonary mucormycosis (20). Our data highly suggests that activation of the EGFR occurs by binding of the fungus to integrin β1 (**Fig. 10B**). Specifically, the use of anti-integrin β1 antibody prevents the *R. delemar-*induced activation of EGFR.

We previously reported on protecting DKA mice from mucormycosis by using antibodies targeting GRP78/CotH3 interactions (14, 15, 27). In these studies, mice were partially protected when the antibodies were introduced alone and maximal protection occurred when anti-CotH3 antibodies were combined with antifungal agents (27), indicating the potential translational benefit of this therapeutic approach. In this study, we also demonstrated partial but highly significant protection against pulmonary mucormycosis when a single administration of anti-integrin β1 is used. This antibody dose translates into ∼ 4.0 mg/kg, which is within the antibody doses currently in clinical practice of 1-15 mg/kg, thereby emphasizing the clinical applicability of this approach. One caveat of an immunotherapeutic approach targeting host cell receptors such as integrins or GRP78 is the potential host toxicity. However, it is prudent to point that targets such as GRP78, integrins, or EGFR are the subject of developing and/or developed therapeutic strategies against cancer (36, 37). One advantage of developing therapies targeting integrins would be the possibility of using the developed therapy to treat aspergillosis since we showed that integrin β1 was also identified as a host receptor on A549 alveolar epithelial cells when interaction with *A. fumigatus* cell surface protein CalA (25). *A. fumigatus* CalA specifically interacts with integrin α5β1 subunit, rather that integrin α3β1, the predominant receptor for *R. delemar*. However, blocking of integrin α5 or α5β1 also resulted in a modest yet detectable decrease in *Rhizopus* invasion of alveolar epithelial cells, indicating that the α5 subunit may play a minor role in fungal interactions. Therefore, a therapy that targets both infections would have to focus on targeting integrin β1. Finally, nasal and/or alveolar epithelial cell interactions are early steps of the disease, and any potential therapy targeting these interactions are likely to be more successful if initiated early on, preferably with antifungal therapy to block invasion and enhance fungal clearance. Unfortunately, diagnosis of mucormycosis often occurs in late-stage disease and currently reliant on histopathology or non-specific radiological methods (38). However, early results of several methods reliant on molecular diagnosis (including those targeting CotH genes (39-42)) and serology (targeting mannans (43)) are encouraging and likely to help in implementing early therapy.

To summarize, the unique susceptibility of DKA subjects to rhinocerebral mucormycosis is likely due to specific interaction between nasal epithelial cell GRP78 and fungal CotH3, the expression of which increase in the presence of environmental factors present in DKA, which results in trapping inhaled spores in the nasal cavity. In contrast, pulmonary mucormycosis is initiated via interaction of inhaled spores expressing CotH7 with integrin α3β1 receptor which activates EGFR to induce fungal invasion of host cells. These results add to our previously published line of evidence on the pathogenesis of mucormycosis in different hosts, and provide groundwork for the development of therapeutic interventions against lethal drug-resistant mucormycosis.

## Methods

### *R. delemar* and culture conditions

A variety of clinical Mucorales isolates was used in this study. *R. delemar* 99-880 (brain isolate from a patient with rhinocerebral mucormycosis), *R. oryzae* 99-892 (isolated from a patient with pulmonary mucromycosis) and *Mucor circinelloides* 131 were obtained from the Fungus Testing laboratories at University of Texas Health Science Center at San Antonio (UTHSCSA), Texas. *Lichtheimia corymbifera* strain 008-0490 and *Rhizomucor* were collected from patients enrolled in the The Deferasirox-AmBisome Therapy for Mucormycosis study (DEFEAT Mucormycosis) (44), *Cunninghamella bertholletiae* 182 is a clinical isolate obtained from Dr. Thomas Walsh, (NIH, Bethesda, Maryland, USA). *Saccharomyces cerevisiae* ATCC 62956 (LL-20), its his3Δ and leuΔ, was constructed by L. Lau (University of Illinois at Chicago). *S. cerevisiae* expressing *R. delemar* CotH3 protein driven by the galactose inducible promoter (15) was utilized to confirm the candidate ligand for the nasal epithelial cells. Mucorales were grown on PDA plates (BD Biosciences — Diagnostic Systems) plates for 3–5 days at 37°C, while *S. cerevisiae* was grown on synthetic dextrose minimal medium (SD) for 3-5 days. All incubations were done at 37°C. To induce the expression of CotH3 in *S. cerevisiae*, the yeast cells were grown in synthetic galactose minimal medium (SG) at 37°C for 16 hours. The sporangiospores were collected in endotoxin-free Dulbecco’s phosphate buffered saline (PBS) containing 0.01% Tween 80 for Mucorales, washed with PBS, and counted with a hemocytometer to prepare the final inoculum. For *S. cerevisiae*, cells were centrifuged, and washed with PBS and counted as above.

To form germlings, spores were incubated in Kaighn’s Modification of Ham’s F-12 Medium (F-12K from the American Type Culture Collection [ATCC]) medium at 37°C with shaking for 1–3 hours based on the assay under study. Germlings were washed twice with F-12 Medium for all assays used, except in experiments involving isolation of the epithelial cell receptor, for which the germlings were washed twice with PBS (plus Ca^++^ and Mg^++^).

### Host cells

Nasal epithelial cells (CCL-30) were obtained from ATCC and cultured in Eagle’s Minimum Essential Medium (EMEM) supplemented with 10% fetal bovine serum and penicillin-streptomycin. Homo sapiens alveolar epithelial cells (A549 cells) procured from ATCC were obtained from a 58-year-old male Caucasian patient with carcinoma. They were propagated in F-12 Medium developed for alveolar A549 epithelial cells. The GD25 and β1AGD25 cell lines were obtained from Dr. Deane F. Mosher, University of Wisconsin-Madison. The cells were cultured to confluency in Falcon Tissue Culture Treated Flasks (75cm^2^) at 37°C with 5% CO2.

### Invasion of *R. delemar* to epithelial cells

The number of organisms invading epithelial cells was determined using a modification of our previously described differential fluorescence assay (45). Briefly, 12-mm glass coverslips in 24-well cell culture plate were coated with fibronectin for at least 4 hours and seeded with epithelial cells until confluency. After washing twice with prewarmed Hank’s Balanced Salt Solution (HBSS, Irvine Scientific), the cells were then infected with 2.5 × 10^5^ cells of *R. delemar* in F-12K medium that had been germinated for 2 hours. Following incubation for 3 hours, the cells were fixed in 3% paraformaldehyde and were stained for 1 hour with 1% Uvitex (Polysciences), which specifically binds to the chitin of the fungal cell wall. After washing 5 times with PBS, the coverslips were mounted on a glass slide with a drop of ProLong Gold antifade reagent and sealed with nail polish. The total number of cell-associated organisms (i.e., germlings adhering to monolayer) was determined by phase-contrast microscopy. The same field was examined by epifluorescence microscopy, and the number of uninternalized germlings (which were brightly fluorescent) was determined. The number of endocytosed organisms was calculated by subtracting the number of fluorescent organisms from the total number of visible organisms. At least 100 organisms were counted in 20 different fields on each slide. Two slides per arm were used for each experiment, and the experiment was performed in triplicate on different days.

### *R. delemar* –induced epithelial cell damage

Host cell damage was quantified by using a chromium (^51^Cr)-release assay (46). Briefly, epithelial cells grown in 24-well tissue culture plates were incubated with 1 μCi per well of Na_*2*_^51^CrO_4_ (ICN) in EMEM or F12-K medium (for nasal or alveolar cells) for 16 hours. On the day of the experiment, the unincorporated ^51^Cr was aspirated, and the wells were washed twice with warmed HBSS. Cells were infected with 2.5 × 10^5^ spores suspended in 1 ml in EMEM or F-12K. Spontaneous ^51^Cr-release was determined by incubating epithelial cells in EMEM or F-12K medium without *R. delemar.* After 30 hours of incubation of spores with nasal cells, or 48 hours for alveolar cells, 50% of the medium was aspirated from each well and transferred to glass tubes. Approximately 500 µl of 6 N NaOH was added to each well, incubated for 15 min, and the media transferred from the wells to a glass tube. Subsequently each well was rinsed with 500 µl of RadiacWash (Biodex), and transfer to the same tube. The amount of ^51^Cr in the tubes was determined by gamma counting. The total amount of ^51^Cr incorporated by epithelial cells in each well equaled the sum of radioactive counts per minute of the aspirated medium plus the radioactive counts of the corresponding cells. After the data were corrected for variations in the amount of tracer incorporated in each well, the percentage of specific epithelial cell release of ^51^Cr was calculated by the following formula: [(experimental release × 2) – (spontaneous release × 2)]/[total incorporation – (spontaneous release × 2)]. Each experimental condition was tested at least in triplicate and the experiment repeated at least once.

For Ab-mediated blocking of adherence, invasion or damage caused by *R. delemar*, the assays were carried out as above except that epithelial cells were incubated with the respective antibodies [50 μg/ml anti-GRP78 or 5 μg/ml anti-integrin β1 or integrin α3β1 Ab or Anti-IgG (as an isotype matching control)] for 1 hour prior to addition of *R. delemar* germlings.

### Effect of acidosis, iron, glucose or β-hydroxy butyrate on *R. delemar* –epithelial cell interactions

Studies were performed to investigate the effect of glucose, iron or BHB on epithelial cell GRP78 or integrin expression levels, and to test their impact on subsequent interactions of epithelial cells with *R. delemar* germlings. Epithelial cells were grown in EMEM or F-12K media containing varying concentrations of FeCl_3_, glucose or BHB for 5 hours. GRP78 or integrin expression, invasion, and damage assays were conducted as mentioned in the previous section.

### Extraction of epithelial cell membrane proteins

Epithelial cell membrane proteins were extracted according to the method of Isberg and Leong (21). Briefly, epithelial cells grown to confluency in 20 flasks of 75 cm^2^ were split into ten tissue culture dishes 150 mm × 25 mm and incubated at 37°C in 5% CO_2_ until they reached confluency (normally 5-7 days). The cells were washed two times with 12 ml warm PBS containing Ca^++^ and Mg^++^ (PBS-CM) prior to incubating them with 0.5 mg/ml EZ-Link sulfo-NHS-LS-Biotin (Pierce) (12 minutes in 5% CO_2_ at 37°C). Subsequently, the cells were then rinsed extensively with cold PBS-CM and scraped from the tissue culture dishes. The epithelial cells were collected by centrifugation at 500 g for 5 minutes at 4°C and then lysed by incubation for 20 minutes on ice in PBS-CM containing 5.8% n-octyl-β-d-glucopyranoside (Fisher) and protease inhibitor cocktail solution (Fisher). The cell debris was removed by centrifugation at 5,000 g for 5 minutes at 4°C. The supernatant was collected and centrifuged at 100,000 g for 1 hour at 4°C. The concentration of the epithelial cell proteins in the resulting supernatant was determined using the Bradford method (Bio-Rad).

### Isolation of epithelial cell receptors that bind to Mucorales

Live Mucorales spores (1 × 10^8^) or an equivalent volume of 1–3 hours germlings (approximately 1 × 10^8^ cells) were incubated for 1 hour on ice with 250 μg of biotin-labeled epithelial cell surface proteins in PBS-CM plus 1.5% n-octyl-β-d-glucopyranoside and protease inhibitor cocktail. The unbound epithelial cell proteins were washed away by 5 rinses with this buffer. The epithelial cell proteins that remained bound to the fungal cells were eluted twice with 6 M urea (Sigma). The proteins were then separated on 10% SDS-PAGE and transferred to immun-Blot PVDF Membrane (BIO-RAD). The membrane was then treated with Western Blocking Reagent (Roche) and probed with anti-biotin, HRP conjugated linked antibody (Cell Signaling). After incubation with SuperSignal West Dura Extended Duration Substrate (Pierce), the signals were detected using a CCD camera. To identify epithelial cell proteins that bound to Mucorales, we incubated epithelial cell membrane proteins with *R. delemar* germlings as above. The eluted proteins were separated by SDS-PAGE, and the gel was stained with Instant Blue Stain (Fisher). The major two bands at approximately 75 and 130 kDa (from alveolar and nasal cells) were excised and micro sequenced using MALDI-TOF MS/MS (The Lundquist Institute Core Facility).

To confirm the identity of GRP78 and integrin α3β1, epithelial cells membrane proteins that bound to *R. delemar* were separated on an SDS-polyacrylamide gel and transferred to PVDF-plus membranes. Membranes were probed with a rabbit anti-GRP78 antibody (Abcam), followed by HRP-conjugated goat anti-rabbit IgG (Pierce) as a secondary Ab (for nasal cells) and rabbit anti-integrin α3β1 (Abcam), followed by HRP-conjugated goat anti-rabbit IgG (Pierce). After incubation with SuperSignal West Dura Extended Duration Substrate (Pierce), the signals were detected using enhanced chemiluminescence and imaged with a C400 (Azure Biosystems) digital imager.

### Immunoblot of EGFR phosphorylation *in vitro*

A549 cells in 24-well tissue culture plates were incubated in F-12K tissue culture medium supplemented with fetal bovine serum to a final concentration of 10%. Prior to infection, the A549 cells were serum starved for 120 minutes. Spores of *R. delemar* was incubated in RPMI for 60 minutes at 37°C, washed, and suspended in F-12K medium. A549 cells were infected for 3 hours with a multiplicity of infection (MOI) of 5. Next, the cells were rinsed with cold HBSS containing protease and phosphatase inhibitors and removed from the plate with a cell scraper. After collecting the cells by centrifugation, they were boiled in 2x SDS sample buffer. The lysates were separated by SDS-PAGE, and Y1068 EGFR phosphorylation was detected with a phospho-specific antibody (Cell Signaling). The blots were then stripped, and total protein levels was detected by immunoblotting with appropriate antibodies against EGFR (Cell Signaling). The immunoblots were developed using enhanced chemiluminescence and imaged with a C400 (Azure Biosystems) digital imager.

### Colocalization of GRP78 and integrin α3β1 with phagocytosed *R. delemar* germlings

We used a modification of our previously described method (14). Confluent epithelial cells on a 12-mm-diameter glass coverslip were infected with 2.5 × 10^5^ cells/ml *R. delemar* cells in EMEM or F12-K mediums that had been pregerminated for 2 hours. After 3 hours incubation at 37°C, the cells were gently washed twice with HBSS to remove unbound organisms, and then fixed with 3% paraformaldehyde for 15 min.

For *R. delemar* interaction with nasal cells, a proximity ligation assay technique (PLA, Sigma Aldrich) was performed. For the PLA assay, two primary antibodies raised in different species are used to detect two unique protein targets. A pair of oligonucleotide-labeled secondary antibodies (PLA probes) then binds to the primary antibodies. Hybridizing connector oligos join the PLA probes only if they are in close proximity to each other, allowing for up to 1000-fold amplified signal tethered to the PLA probe, resulting in localization of the signal. This is visualized and quantified as discrete spots (PLA signals) by microscopy image analysis. Thus, two different antibodies - a mouse anti-GRP78 IgG was used to stain paraformaldehyde-fixed nasal cells, while anti-rabbit IgG against CotH3 was used to label *R. delemar*. Interaction between the two cell-surface proteins were carried out according to the kit instructions and visualized by confocal microscopy.

For alveolar epithelial cell-*R. delemar* interaction, the formaldehyde-fixed epithelial cell-spore mixture were incubated with 1% BSA for 1 h (blocking step). Next, cells were incubated with antibodies against integrin α3 or integrin β1 (eBioscience, Santa Cruz), followed by the appropriate secondary antibodies labeled with either Alexa Fluor 488 or Alexa Fluor 568 (Thermo Fisher Scientific). After washing, the coverslip was mounted on a glass slide with a drop of ProLong Gold antifade reagent (Molecular Probes, Invitrogen) and viewed by confocal microscopy. The final confocal images were produced by combining optical sections taken through the z axis.

### Protoplast formation and collection of *R. delemar* cell wall material

To identify the *R. delemar* ligand that binds to epithelial cell GRP78, we collected cell wall material from supernatants of protoplasts of *R. delemar* germlings. Briefly, *R. delemar* spores (6 × 10^6^) were germinated in YPD medium for 3 hours at 37°C. Germinated cells were collected by centrifugation at 900 g, washed twice with 0.5 M sorbitol, and then resuspended in 0.5 M sorbitol in sodium phosphate buffer (pH 6.4). Protoplasting solution consisting of 0.25 mg/ml lysing enzymes (Sigma-Aldrich), 0.15 mg/ml chitnase (Sigma-Aldrich), and 0.01 mg/ml chitosinase (produced from *Bacillus circulans*) was added to the germinated spores and incubated with gentle shaking at 30°C for 2 hours. Protoplasts were collected by centrifugation for 5 minutes at 200 g at 4°C, washed twice with 0.5 M sorbitol, and resuspended in the same buffer. Incubating protoplasts in the presence of the osmotic stabilizer sorbitol enables regeneration of the cell wall, and during regeneration, cell wall constituents are released into the supernatant (47-49), protoplasts were pelleted, and the supernatant was sterilized by filtration (0.22-μm filters) in the presence of protease inhibitors (Pierce). The supernatant was concentrated, and protein concentration was measured using the Bradford method (BioRad). Negative control samples were processed similarly, with the exception of the absence of protoplasts. Thus, far-western blot analysis using recombinant human GRP78 and anti-GRP78 antibodies was done to identify *R.delemar* ligand.

### *In vivo* virulence studies

For survival studies, equal numbers of male and female ICR mice (≥20 g) were purchased from Envigo and housed in groups of 5 each. Mice were immunosuppressed with cyclophosphamide (200 mg/kg i.p.) and cortisone acetate (500 mg/kg s.c.) on day -2, +3, and +8. Mice were infected with 2.5 × 10^5^ in 25 μl *R. delemar* spores intratracheally. To confirm the inoculum, 3 mice were sacrificed immediately after inoculation, their lungs were homogenized in PBS and quantitatively cultured on PDA plates containing 0.1% triton, and colonies were counted after a 24-hour incubation period at 37°C. Mice were treated with a single dose of 100 μg (i.p.) anti-β1 integrin antibody administrated 24 h post infection. Placebo mice received 100 µg of isotype-matched IgG. Mouse survival was monitored for 21 days, and any moribund mice were euthanized. Results were plotted using Log-rank (Mantel-Cox) Test.

### Study approval

All procedures involving mice were approved by the IACUC of The Lundquist Institute for Biomedical Innovations at Harbor-UCLA Medical Center, according to the NIH guidelines for animal housing and care. Human endothelial cell collection was approved by the IRB of The Lundquist Institute for Biomedical Innovations at Harbor-UCLA Medical Center. Because umbilical cords are collected without donor identifiers, the IRB considers them medical waste not subject to informed consent.

### Statistical analysis

Differences in GRP78 or integrin β1 expression and fungi–epithelial cell interactions were compared by the nonparametric Mann-Whitney test. In the survival study, the nonparametric log-rank test was used to determine differences between isotype IgG control and Ant-integrin β1 Ab. Comparisons with *P* values < 0.05 were considered significant.

## Acknowledgments

This work was supported by a Public Health Service grant R01AI063503, R01AI141360 and SBIR 5R43AI138904 to ASI. MS is supported by R00DE026856, VMB by U19AI110820 and 358 R01AI141360, PU by 1R21HD097480-01 and R01AI141794 and SGF by R01AI124566 and R01DE022600.

We would like to thank Drs. H. K. Choi, Davood Soleymani, and Helen Chun for their guidance and helpful discussions.

## Supplemental Material

**Fig. S1. *R. delemar* damage of primary human alveolar epithelial cells (PAEpiC).** Damage assay time points were carried out using ^51^Cr release method. Data are expressed as median ± interquartile range.

**Fig. S2. Expression of GRP78 and integrin during *R. delemar* infection of alveolar epithelial cells after 3 hours of interaction.** The expression was quantified by qRT-PCR. Data are expressed as median ± interquartile range from three independent experiments.

**Fig. S3. Co-localization of integrin β1 and α5 around *R. delemar***: Confocal microscopy images showing differentially fluorescent integrin β1 but not α5 around *R. delemar* germlings during infection of alveolar epithelial cells. Images were taken after 2.5 hours of incubation of the fungus with the host cells.

**Fig. S4. RNAi targeting CotH7 inhibits the expression of CotH7.** *R. delemar* was transformed with an RNAi construct targeting CotH7 expression or empty plasmid. Cells transformed with RNAi construct targeting CotH7 demonstrated 50% reduction in CotH7 expression relative to empty plasmid-transformed *R. delemar*, as determined by RT-PCR after 16 hours of incubation.

**Fig. S5. Inhibition of CotH3 had no effect on invasion of or damage to alveolar epithelial cells by *R. delemar***. Inhibition of CotH3 expression by RNAi did not alter the ability of *R. delemar* to adhere to, invade or damage alveolar epithelial cells vs. *R. delemar* transformed with empty plasmid (A). Anti-CotH3 antibody block *R. delemar* mediated adhesion and invasion but not alter damage of alveolar cells when compared to host cells incubated with isotype-matched IgG (B). Adhesion and invasion assay was carried out by differential fluorescence, while damage was carried out using ^51^Cr release method. Data are expressed as median ± interquartile range from 3 independent experiments.

**Fig. S6. Inhibition of CotH7 by RNAi had no effect on *R. delemar* interactions with nasal epithelial cells.** Adhesion and invasion assay was conducted by differential fluorescence using nasal cells spilt on 12-mm glass coverslips, while damage assay was carried out using ^51^Cr release assay. Data are expressed as median ± interquartile range from 3 independent experiment.

**Fig. S7. The CotH protein family.** Phylogenetic tree and relative distance of *R. delemar* CotH proteins (A), and their percent identity (B).

**Fig. S8. Alignment results between CotH3 peptide (that has been used for anti-CotH3 production) and CotH7.** Multiple Sequence Comparison by Log-Expectation (MUSCLE) online tool used to perfume sequence alignment between 16-mer CotH3 and CotH7 protein using cluster 12.1 algorithm.

